# Characterization of yield and fruit quality parameters of Vietnamese elite tomato lines generated through phenotypic selection and conventional breeding methods

**DOI:** 10.1101/2022.11.28.518131

**Authors:** Cam Chau Nguyen, Rahul Mahadev Shelake, Tien Vu Van, Hai Tong Van, Nhan Nguyen Thi, Xuan Canh Nguyen, Vo-Anh-Khoa Do, Hai Nguyen Thanh, Jae-Yean Kim

**Author notes:** Correspondence: Cam Chau Nguyen, Jae-Yean Kim. Co-authors: Rahul Mahadev Shelake Tien Vu Van Hai Tong Van Nhan Nguyen Thi Xuan Canh Nguyen Vo-Anh-Khoa Do Hai Nguyen Thanh.

## Abstract

Tomato (*Solanum lycopersicum* L.) is the second most important vegetable crop after potatoes, and global demands have been steadily increasing in recent years. Conventional breeding has been applied to breed and domesticate tomato varieties to meet the need for higher yield or superior agronomical traits that allow to sustain under different climatic conditions. In the current study, we applied bulk population breeding by crossing eight tomato accessions procured from the Asian Vegetable Research and Development Center (AVRDC) with three heat-resistant tomato inbred lines from Vietnam and generated ten elite tomato (ET) lines in the F8 generation. The individual F8 lines exhibited robust vigor and adaptability to Vietnamese climate conditions. Among the ten lines, ET1 and ET3 displayed indeterminate growth. ET2 showed semi-determinate, while all the other lines had determinate growth. The different ET lines showed distinctive superior agronomical traits, including early maturing (ET4, ET7, and ET10), highly efficient fruit set (ET1), higher yield (ET1, ET8, ET10), jointless pedicels (ET2), and partial parthenocarpy (ET9). Molecular analysis revealed that the ET3 line consisted of *Ty-1* and *Ty-3* loci that positively contribute to *Tomato yellow leaf curl virus* (TYCLV) resistance in tomato plants. The elite tomato lines developed in this study would contribute significantly to the Vietnamese and Asian gene pool for improved tomato production and may be a valuable resource for various breeding goals.

## 1 Introduction

Tomato breeding has come a long way towards turning wild tomato relatives into commercial cultivars, with a vast number of varieties marketed across the globe (Robertson and Labate 2006; Bai and Lindhout 2007). All the domesticated tomato belongs to *Solanum lycopersicum*, one of 3000 species of the Solanaceae family (Knapp 2002). Conventional breeding methods have played a pivotal role in the development and domestication of important elite tomato varieties with outstanding agronomic traits (Cappetta et al. 2020). Seed companies compete fiercely to gain a part of the seed market, which is about $1.09 billion worldwide (Research and Markets 2022). The major tomato seed companies are located in North America, while some other companies are in Asia Pacific, which is expected to be the fastest-growing market. Farmer- or consumer-favorable traits are the major determinants while developing new varieties, which include higher yield, high-quality fruits, and biotic and abiotic stress tolerance. China and India are the leading markets and producers of tomatoes not only in Asia but also in the world, which produced 64.87 million tons and 20.57 million tons in 2020, respectively (FAO 2020). Asian countries, therefore, need to import a large amount of tomato seeds from American companies. The unequal needs and supply sources of tomato seeds in Asia require fast-growing seed providers with the best quality traits.

In the domestication process of new cultivars with superior characteristics, conventional breeding takes advantage of the crossing techniques, which is favorable to incorporating desired traits without affecting the natural boundaries of an organism (Acquaah 2012). Historically, traditional breeding is mainly based on the morphological discrimination/selection of plants in the previous season to collect the material seeds for the following year. These superior plants provide a rich source of genetic diversity for the next generation, although phenotypes may be quite uniform. Modern conventional breeding requires scientific background about the target cultivars and a keen sense of observation (Tester and Langridge 2010). Two processes are often involved in classic breeding: variability assembling and discrimination to select and improve the desired traits (JUSTIN and Fehr 1988). Associated techniques exploited to facilitate more effective conventional breeding include artificial pollinations, hybridization, wild crosses, embryo culture, chromosome doubling, doubled haploids, bridge crossing, protoplast fusion, seedlessness, genetic marker, and mutagenesis (Acquaah 2016).

Research regarding tomato improvement in terms of productivity and quality is in high demand to attain higher yields and satisfy industrial or consumer requirements. One of the most important genetic sources of tomato, as well as other vegetables, is the World Vegetable Center, which originated from the Asian Vegetable Research and Development Center (AVRDC) founded in 1971. Located in Asia and Africa, every year, this largest vegetable gene bank provides about 10,000 seed samples belonging to 65,152 accessions of 330 species from 155 countries for agronomical research all around the world.

The desirable traits that are considered to be the objectives of breeding are dependent on the demands of end-users (Acquaah 2016). Some examples include plant architecture, flowering and fruit ripening time, fruit shape and weight, fruit number, productivity, sweetness, and advantageous traits such as jointless and seedless fruits. Considering the demands of a large population in Asian countries, high productivity is the major goal of most tomato breeding programs. Yield performance is mainly determined by fruit set efficiency and fruit size, which are analyzed in terms of various parameters, such as fruit number, cluster number, fruit forming ratio, fruit weight, and the number of locules (de Souza et al. 2012). Another favorable trait for tomato growers is early maturity, which is correlated with early flowering and quick ripening. Reduction of growing time benefits the producers by lowering labor cost and risk of exposure to biotic and abiotic damages. Early fruits are often priced higher in the market, thus increasing production efficiency, and are profitable to farmers. One of the factors that determine maturing time is plant growth habit/architecture. Tomato stem architecture includes determinate, semi-determinate, and indeterminate (Vicente et al. 2015). Finally, fruit quality is the most judged trait by consumers, thereby determining the price of tomato in the market. Depending on the purposes, such as direct eating, cooking, or industrial processing (mainly to produce juice and paste), the requirements of fruit quality are different. Sweetness is the most basic criteria to determine the flavor of tomato. The sweet taste in tomato is affected by the levels and ratios of sugars such as glucose, fructose, and sucrose (Baldwin et al. 2000; Georgelis et al. 2004). Total soluble solid (TSS) include sugar contents that can be measured by the Brix scale, in which one degree equals to 1% of sugar concentration in the solution at 20 °C. The TSS value can be easily obtained by a Brix refractometer, which is very convenient, cost-efficient, and reliable (Pisello et al. 2021). The average Brix values of tomato fruits in the market ranges from 3.5 to up to 10 depending on the cultivar type (Kleinhenz and Bumgarner 2012). Approximately 5 Brix is expected for cooking tomato, while Brix value >8 is desirable for direct eating.

For tomato growers, plant vigor is one of the most important traits that enhances plant ability to resist major diseases and secure the plant yield. One such disease is the yellow leaf curl caused by *Tomato yellow leaf curl virus* (TYCLV), a *Begomovirus* belonging to the family Geminiviridae. Yellow leaf curl is one of the most damaging viral diseases in tomato production worldwide (Prasad et al. 2020). TYLCV infection could result in 20-100% yield loss, hence development of resistance to this virus is one of the urgent goals of tomato breeding (Levy & Lapidot 2008). Resistant lines could be determined and developed using molecular markers linked to *Resistance* (*R*) genes (Barbieri et al. 2010). There are six TYLCV-related quantitative trait loci (QTLs), namely *Ty-1* to *Ty-6*, that have been exploited to develop resistance in tomato. Among them, *Ty-1, Ty-2*, and *Ty-3* have been focused on the recent breeding programs (Yan et al. 2018). *Ty-1* and *Ty-3* have been proven to contribute to the geminivirus tolerance (Verlaan et al. 2013). Plants possessing *Ty-1* or *Ty-3*, or both the loci were reported to exhibit significant tolerance to TYLCV infection. These loci encode for the same member of RNA-dependent RNA polymerase and may function by generating siRNAs which support geminivirus DNA silencing through direct methylation. *Ty1/Ty3* loci are located on chromosome 6 and could be detected by various molecular markers (Michelson et al. 1994; Zamir et al. 1994; Ji and Scott 2005; Yuanfu Ji et al. 2007). *Ty-1* was mapped to be near the TG97 and TG297 markers (Zamir et al. 1994., Chen et al. 2012). Chen and coauthors (Chen et al. 2012) identified the *Ty-1* by CAPS1 marker followed by *Taq1* enzyme digestion, which could distinguish homozygous and heterozygous plants of this gene. *Ty-3* has two introgression, *Ty-3* from LA27797 and *Ty-3a* from LA1932, which could be identified by a SCAR marker proposed by Ji and coworkers (Ji et al. 2007). Genotypes lacking introgression expressed a 320bp band (*ty-1*), while *Ty-3* and *Ty-3a* presented the fragments of 450 bp and 630 bp, respectively.

In this study, we obtained tomato accessions from AVRDC and then used to cross with domestic tomato lines of Vietnam using a conventional breeding method to produce elite tomato with significant good agronomical traits and well adaptability to Vietnamese climate conditions. Segregating populations were scrutinized based on the above-mentioned phenotypic and morphological characteristics. Elite tomato (ET) lines were developed by the application of artificial crossing, discrimination, and genetic marker analysis. Ultimately, F8 segregated lines were evaluated for plant growth trait, fruit quality, productivity, and beneficial agronomical phenotypes. Also, *Ty-1* and *Ty-3* were assessed in the obtained ET lines. The developed lines would contribute significantly to Vietnamese tomato breeding by expanding the diversity of available gene pool and providing a source of excellent breeding materials.

## 2 Materials and Methods

### 2.1. Plant materials and growth conditions

Ten tomato lines were bred and evaluated following the timeline showed in **Figure 1**. Vietnamese in-bred varieties were selected based on heat resistance and good productivity traits and used as mother lines for breeding. Father lines were obtained from Asian Vegetable Research and Development Center (AVRDC) (**Table 1, Supplementary Table S1**). F8 plants were grown on winter season of 2020 in green house of Vietnam National University of Agriculture (VNUA). The space between plants and between lines was 0.5 meters. At least three plants of each line were used for phenotypic and genomic evaluation. C155, an inbred tomato variety that has been characterized in previous studies and grown widely in Vietnam, was used as a control in this study (Doan et al. 2021; Nguyen et al. 2007).

**Table 1.** Tomato lines developed in this study. Female H: Vietnamese domestic, heat-resistance lines.

| Elite tomato lines<br>(F8) | Parental lines |  | Observed phenotypes |
| --- | --- | --- | --- |
|  | Female | Male |  |
| **ET1** | NHG 195<sup>1</sup> | NHG 176 | high yield |
| **ET2** | H30<sup>2</sup> | NHG 188 | High yield, heat resistance |
| **ET3** | H12 | NHG 191 | High yield, heat resistance |
| **ET4** | H12 | NHG 195 | High yield, heat resistance |
| **ET5** | H30 | NHG 135 | High yield, heat resistance |
| **ET6** | H12 | NHG 188 | High yield, heat resistance |
| **ET7** | H12 | NHG 31 | High yield, heat resistance |
| **ET8** | Similar with ET1 |  |  |
| **ET9** | Similar with ET4 |  |  |
| **ET10** | H67 | NHG 192 | high yield |

<sup>1</sup> NHG lines indicated accessions obtained from AVRDC.
<sup>2</sup> H lines indicated Vietnamese domestic lines which showed heat resistance.

**Figure 1:**
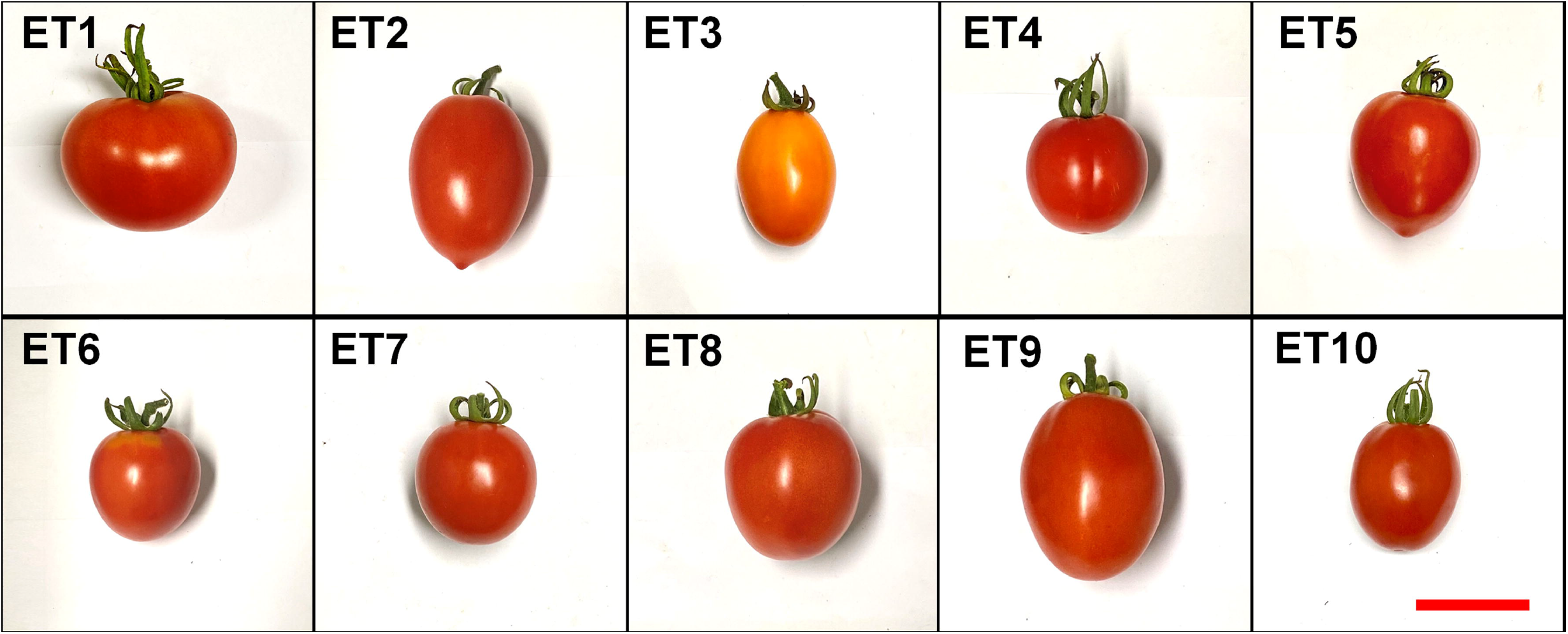
Bulk population breeding of ET lines. Parentals were crossed in 2020. Seeds were continuously harvested and sowed in bulk, followed by selection based on morphological observations. 10% of the best individual plants were selected to grow the next generation until F7. In F7, homogenized plants with beneficial traits were separated to produce F8. Red dots indicate individuals possessing target traits, while white dots indicate plants that do not possess beneficial traits.

### 2.2. Bulk population breeding

Male and female parental varieties were crossed using emasculating and pollinating methods as summarized in earlier study (Markova et al. 2016). Bulk population breeding was performed following the method of (Acquaah 2012). Good fruits of progenies from F1 to F6 were chosen and continuously generated until the F7 generation. On F7 population, the best individual plants were selected and evaluated to develop F8. F8 lines were named as shown in **Table 1** and were used as materials for this study.

### 2.3. Assessment of plant development traits

Grow pattern was determined based on the various recorded traits observed in each line. Plants that showed a halt in shoot development after a fixed period of growth were classified as determinants. Plants that continuously grown and set flowers for a long period were defined as indeterminants. Semi-determinant plants showed both phenotypes described above in each plant.

First flowering and first harvest day were counted from the day of sowing. The fruit forming ratio was calculated by the number of fruits in each of three clusters per plant divided by the number of flowers from the same cluster. The flower number was counted in three representative clusters per plant, and the average was calculated by all the plant samples of each line.

### 2.4. Fruit quality and productivity measurement

Shape index, fruit weight, number of locule, and brix were observed in ten fruits of each plant; and the average was calculated based on all the tested fruits of each line. Shape index (SI) was determined by the ratio between fruit height and fruit width. The Brix level was measured by a Master refractometer (Atago Co. Ltd., Tokyo, Japan). Theoretical yield was calculated by average productivity multiply to 25,000 plants/ha.

### 2.5. Statistical analysis

The data was collated in Excels and analysis by One-way ANOVA with post-hoc Tukey-Kramer’s test (Tukey 1949; Kramer 1956). Tukey-Kramer’s HDS (honest significant difference) is the analysis method to compare the mean between each pairwise combination of groups with unequal sample size. One-way ANOVA was performed by Microsoft Excel Analysis Tools – *ANOVA: Single Factor* with Alpha = 0.05 (Laverty & Kelly 2019; Quirk 2012). Absolute mean difference of each pair was calculated based on the results of One-way ANOVA analysis. Q critical value was identified by the following formula:

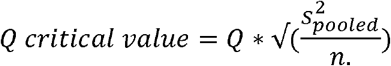

In which, **Q** = Value from Studentized Range Q Table, **s**^**2**^_**pooled**_ **=** Pooled variance across all groups, and **n**. = Sample size for a given group. Finally, the absolute mean difference was compared between each group to identify the Q critical value. The difference between the mean value of two groups is significant if the absolute mean difference value is bigger than Q critical value.

### 2.6. Assessment of TYLCV resistance loci

Genomic DNA was extracted from young leaves of each plant in the investigated population using Qiagen DNeasy Plant Pro Kit (#69204).

Distribution of *Ty-1* locus in ET population was determined by cleaved amplified polymorphic sequence (CAPS) assay (Chen et al. 2012). CAPS1 forward (5’-TAATCCGTCGTTACCTCTCCTT-3’) and reverse (5’-CGGATGACTTCAATAGCAATGA-3’) primers were used to amplify a 398 bp fragment. Afterwards, it was digested by Taq1-v2 (NEB, catalog #R0149S) restriction enzyme to observe 303 bp and 95 bp DNA bands electrophoretically separated on agarose gel.

*Ty-3* was detected by co-dominant sequence characterized amplified region (SCAR) marker P6-25 (Y Ji et al. 2007) which could identify both *Ty-3* (450 bp), *Ty-3a* (630 bp), or *ty-3* (320 bp) alleles. Glyceraldehyde-3-phosphate dehydrogenase (GAPDH) was used as internal control (primers GAPDH-F, 5’-GATTCGGAAGAATTGGCCG-3’ and GAPDH-R, 5’-TCATCATACACACGGTGAC-3’). PCR products were detected by electrophoresis on 1% agarose gel.

## 3 Results

### 3.1. Breeding of elite tomato by crossing between domestic varieties and outstanding cultivars from AVRDC followed by bulk population breeding method

Bulk population strategy was used to exploit natural selection pressure and to improve adaptability of the crossing population from early stage of selection. The individuals possessing less adaptability or low productivity were eliminated from the population. After crossing, about 100 plants of each F1 from a pair of parents were harvested and sowed in bulk (**Figure 1**). Seeds from F2 were harvested and used for planting about 2000 F3 plants. 10% of plants with best viability and productivity were continuously harvested and regenerated until F7 generation. In F7, plants with desirable traits were preliminary selected and separated to produce F8 population. Individual plants of F8 generation were planted in greenhouse and were evaluated by phenotypic and genotypic analyses. Most of the female lines were Vietnamese local lines, while males originated from AVRDC (**Table 1, Supplementary Table S1**). From eight pairs of parents, ten ET lines with superior traits were identified and separately examined (**Table 1**). The developmental and yield-associated traits were assessed in the current work that included the growth pattern, time to first flowering, leaf level to first flowering, flowers per cluster, fruit forming ratio, time to first harvest, and plant height when terminated. Fruit traits (width, height, shape index, weight, number of locules), productivity, and presence of TYLCV-resistance alleles.

### 3.2. Significant good growth and developmental traits were identified in F8 generation

Plant architecture significantly affects the viability and adaptability of tomatoes. It is one of the first factors to be considered when choosing the appropriate cultivars to suit specific cultivation methods and goals (Jo and Shin 2020). Most of examined lines (ET4 to ET10) exhibited determinate growth, only one line (ET2) was semi-determinate, and two lines (ET1, ET3) were indeterminate (**Table 2**). Bushy phenotype of determinate tomato is the favorable trait for growing tomatoes in the open fields. Particularly, Southeast Asian countries, where the parental lines used in this study are originated, prefer the open-field cultivation methods. Although these plants would require a larger space, it helps reduce labor to hold the plants from falling. In addition, bushy plant shape is good for high-productivity varieties since it can carry a large number of fruits.

**Table 2:**
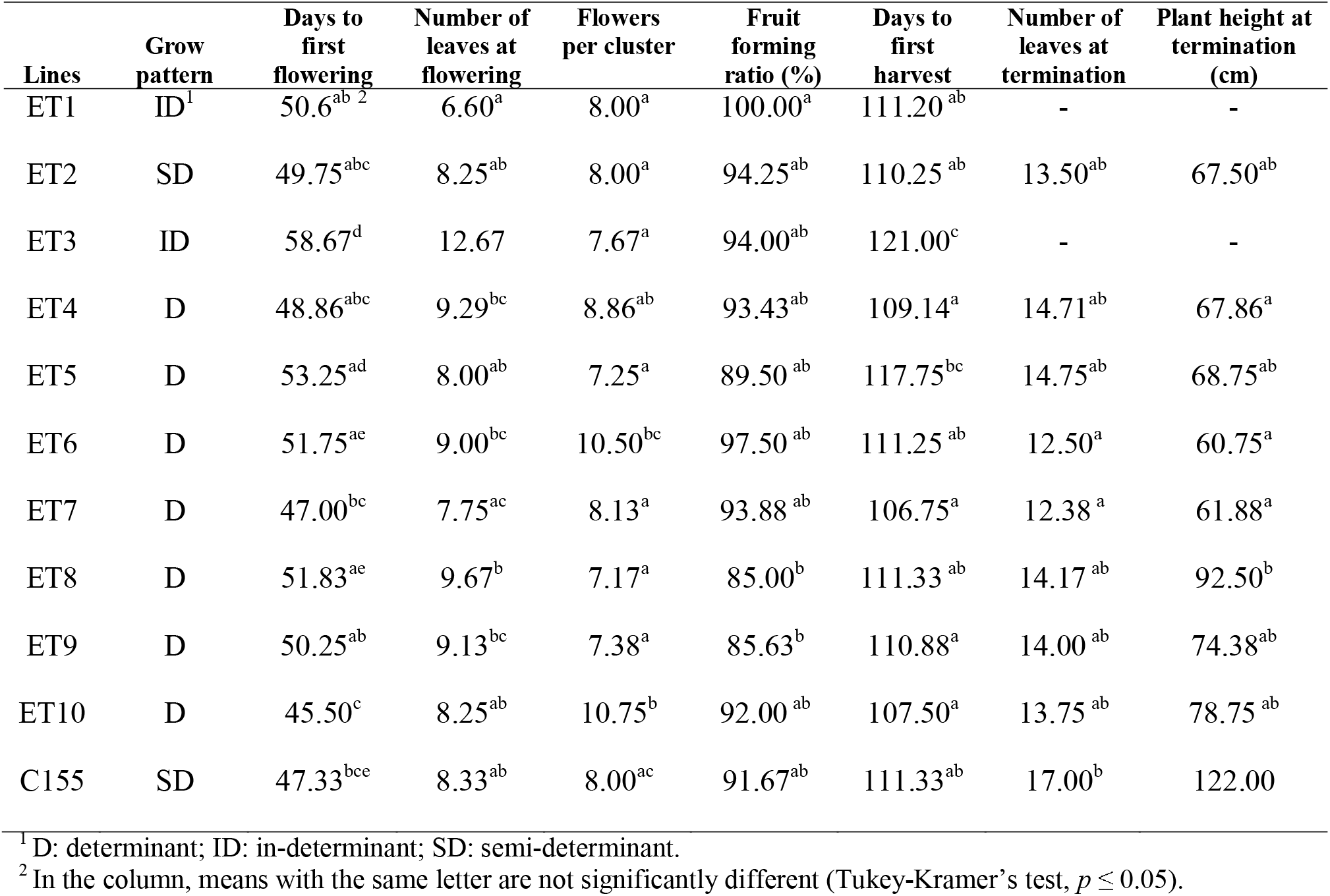
Mean comparison of development and growth traits of F8 plants.

Flowering and fruit-harvesting time are important to determine the length of cultivation. Average flowering time of ET10 (45.5 days post sowing, DPS), ET7 (47 DPS), and ET4 (48.86 DPS) were significantly earlier compared to ET3 (58.67 DPS) (**Table 2**). However, none of the newly developed lines produced flowers earlier than the control C155 line. ET1 showed lowest number of leaves below inflorescence, which was on average 6.6 leaves, indicating the fast development of this line. ET3 produced the first flowers only after having 12.67 leaves on average, which was significantly slower than the control and all the other lines of the population. This phenomenon can be attributed to the nature of indeterminate varieties. Correlated with slow flowering time, ET3 fruits could only be harvested the first time after 121 DPS. Fruits of ET7 were ripened quickly and available for the first harvest at 106.75 DPS. Since the plant height of ET7 line is short – 61.88 cm when terminated, the nutrition might be concentrated to fruit ripening. Thus, ET7 together with ET4 and ET10 (107.5 DPS) could be good candidates for early maturity and quick fruit production. On the other hand, ET3 is significantly slow in flowering and fruit ripening, in comparison with other lines in the population.

Fruit set ratio is important for final production of fruit. Most of ET lines showed good performance regarding fruit forming ratio, except ET8 (85.0%) and ET9 (85.6%), which are significantly lower than the other lines (**Table 2**). Particularly, ET1 expressed 100% fruit forming, which secured the high productivity of this line. Although the number of flowers per cluster of this line (8.0) is not higher than ET2 (8.0), ET4 (8.86) or ET7 (8.13), and significantly less than ET6 (10.5) and ET10 (10.75), best fruit set made ET1 be one of the highest yielding lines in the population, whose total yield is only after ET8 and ET10 (**Table 2, Table 3**).

**Table 3:**
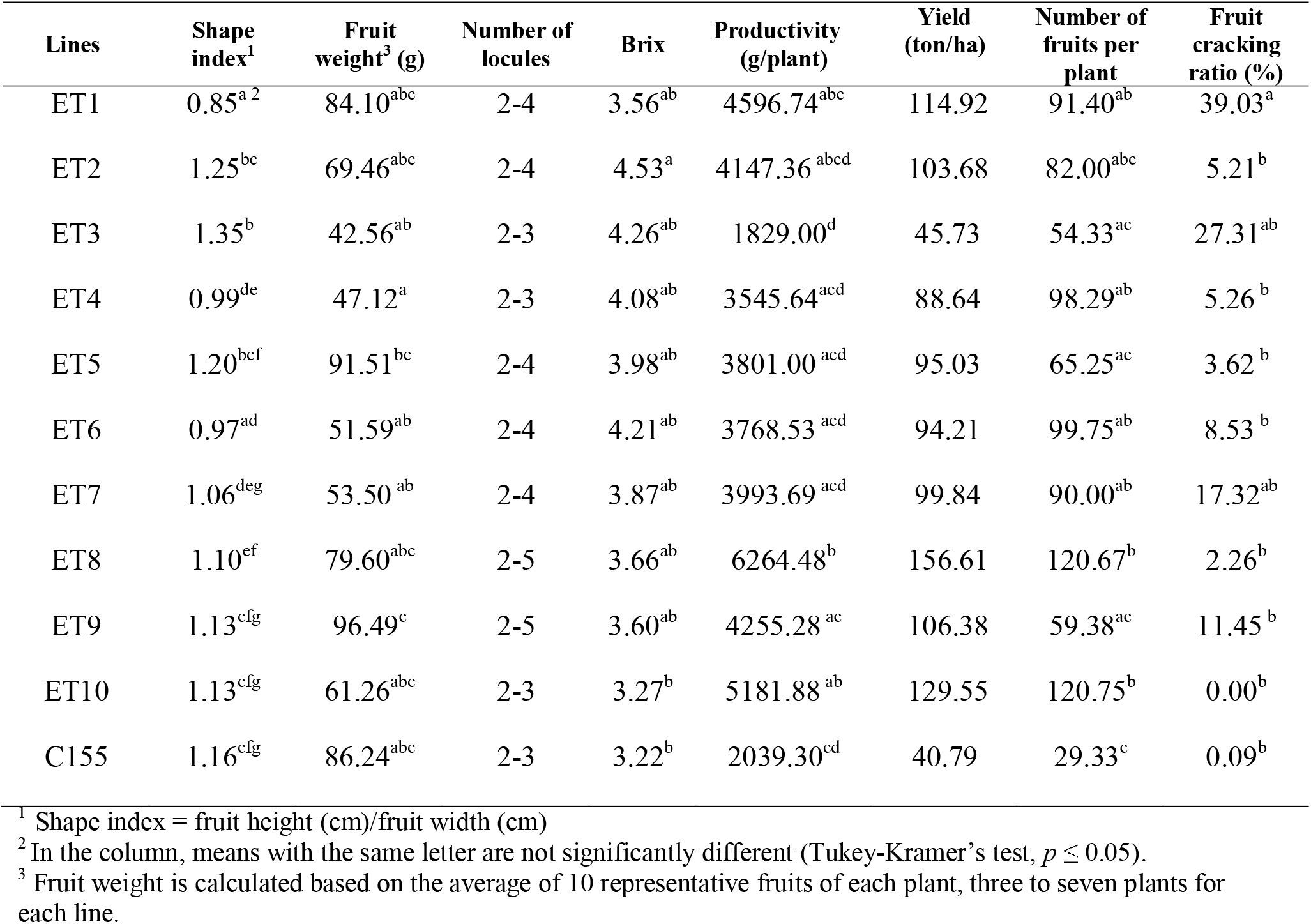
Means comparison of fruit and productivity factors of elite parental lines.

### 3.3. Variable and outstanding traits related to fruit quality and productivity were evaluated in F8 population

The fruit shape and appearance are important for market value because they would affect the preference of customers. Shape index is calculated by fruit length divided to fruit width. The average shape index of F8 lines showed that the fruits of most lines were round-shape, except ET2 and ET3 showed long-shape with a shape index of 1.25 and 1.35, respectively (**Table 3, Figure 2**).

**Figure 2:**
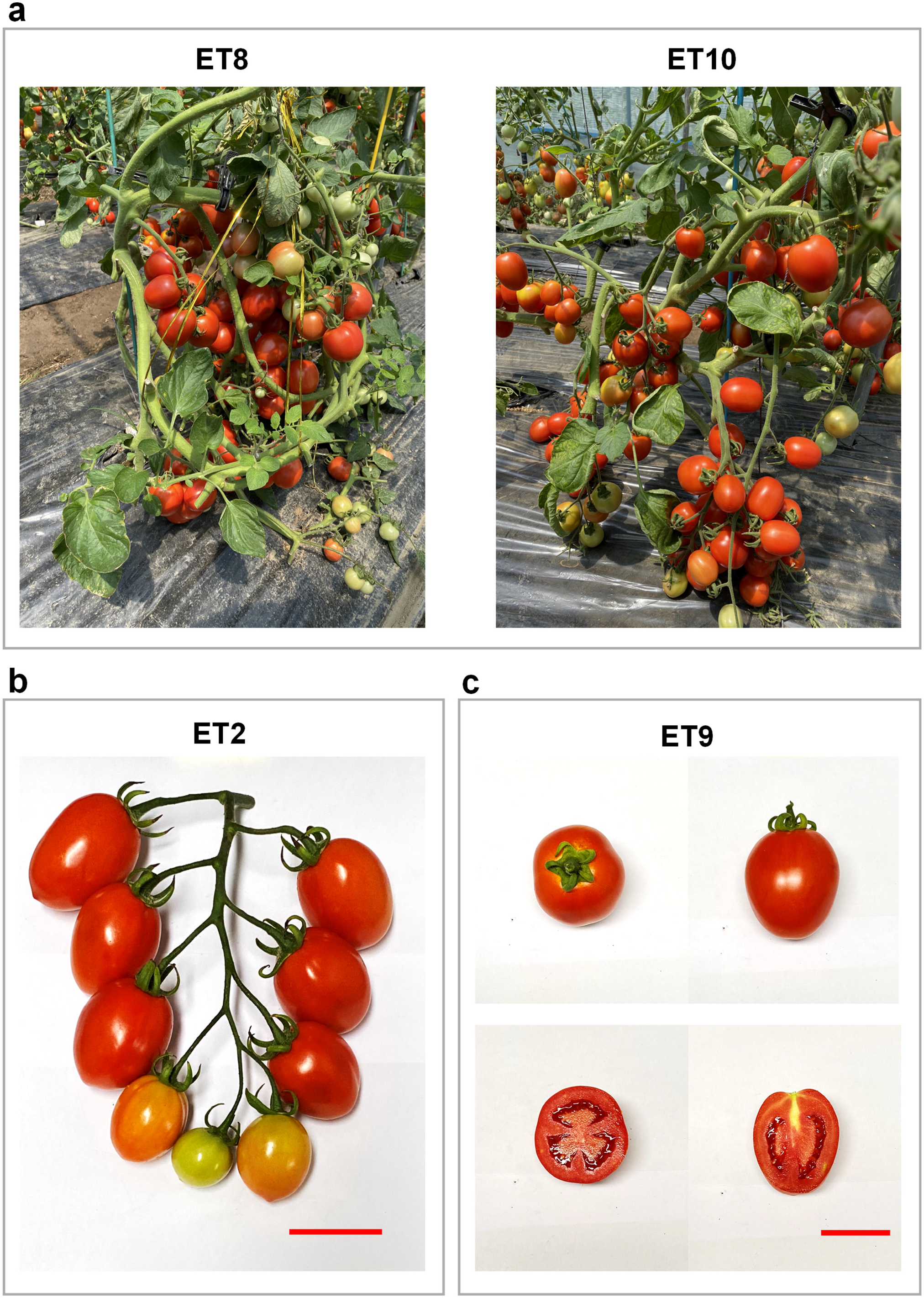
Fruit phenotypes of F8 ET lines. Represent fruits were collected from 100 to 122 days post-sowing depending on the maturation time of each ET line. Bar = 5 cm.

Fruit weight showed a considerable variation, given a wide range of choices for different purposes of tomato production. While ET3 fruits were significantly small, with an average of 42.56 g/fruit, there were lines that produced fruits heavier than 80 g such as ET1 (84.1 g), ET5 (91.51 g), and ET9 (96.49 g) (**Table 3, Figure 2**). Although fruits of ET8 were not bigger compared to other lines, the number of fruits in each plant was the highest (120.75 fruits/plant on average), so the productivity of ET8 was higher than other lines in this study with an average of 6.3 kg/plant (**Table 3, Figure 3a**). Following ET8, ET10 showed a high productivity (5.2 kg/plant) which is also correlated to the fruit number, but not the fruit size. Compared to the control C155, ET8 and ET10 productivity is significantly higher. Correlated with productivity, the theoretical yield of ET8 was the highest with 156.61 ton/ha. The second high yield line was ET10 (129.55 ton/ha), followed by ET1 with 1150 ton/ha. These yielding were more than two to three times of the control, which was 40.79 ton/ha. Therefore, ET1, ET8 and ET10 are good candidates for breeding high yield tomato lines. All the ET lines showed fruit Brix does not excess than 5.0, indicating that these tomato fruits are more suitable for cooking and processing than for eating directly (**Table 3**).

**Figure 3:**
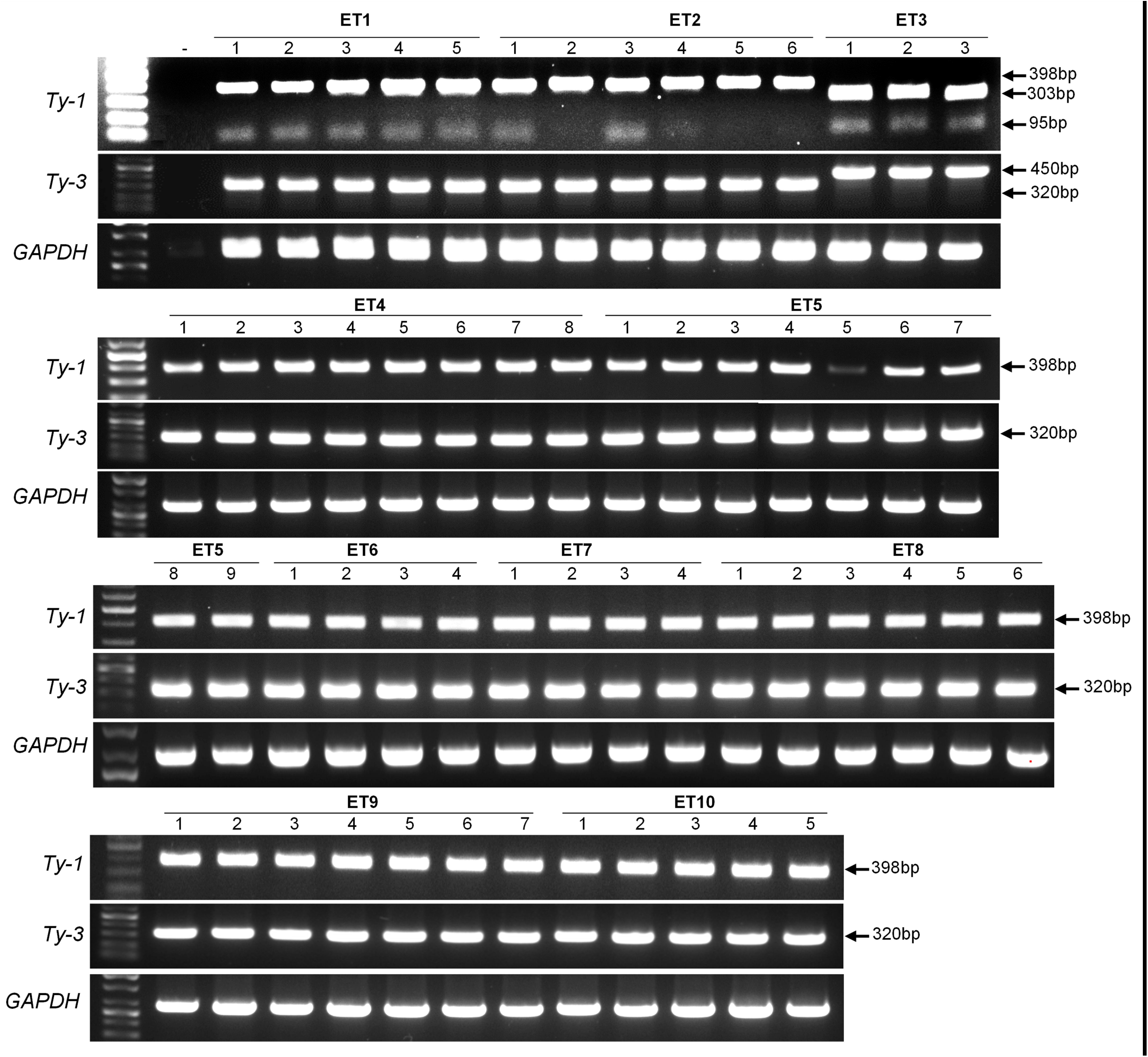
Superior traits of ET lines (F8). a, High-yield lines ET8 and ET10; b, Jointless fruits of ET2; c, Parthenocarpy phenotype of ET9. Bar = 5 cm.

### 3.4. Analysis of specific advantageous phenotypes in F8 generation

Several specific advantageous characteristics were observed in the F8 ET population. For example, ET2 showed a jointless phenotype in its pedicels, which is correlated with low abscission of flowers and fruits (**Figure 3b**). This aspect is of great significance during fruit harvesting since it helps to reduce the loss of flowers or fruits and easily supports the removal of flower stems from fruits. Average fruit forming ratio of ET2 line was 94.3% (**Table 2**), indicating the advantage of jointless phenotype for maintaining fruits on plants.

ET9 expressed partial parthenocarpy phenotype with less than ten seeds per fruit (**Figure 3c**). Together with its big fruit (94.7g/fruit on average), this line provides favorable fruit traits for the market.

### 3.5. TYLCV-resistant alleles in ET lines

For assessment of TYLCV resistance alleles, samples were taken from at least 3 plants of each line. For *Ty-1*, three plants of ET3 line showed 303bp and 95bp bands, indicating that this line possesses *Ty-1* allele while the other lines did not (**Figure 4**). Similarly, only ET3 plants showed 630 bp bands when amplified by SCAR marker P6-25, which is typical for the *Ty3a* allele. This allele assessment denoted that ET3 possesses two TYLCV-resistant alleles, which may contribute to plants’ responses to this dangerous ssDNA begomovirus.

**Figure 4.**
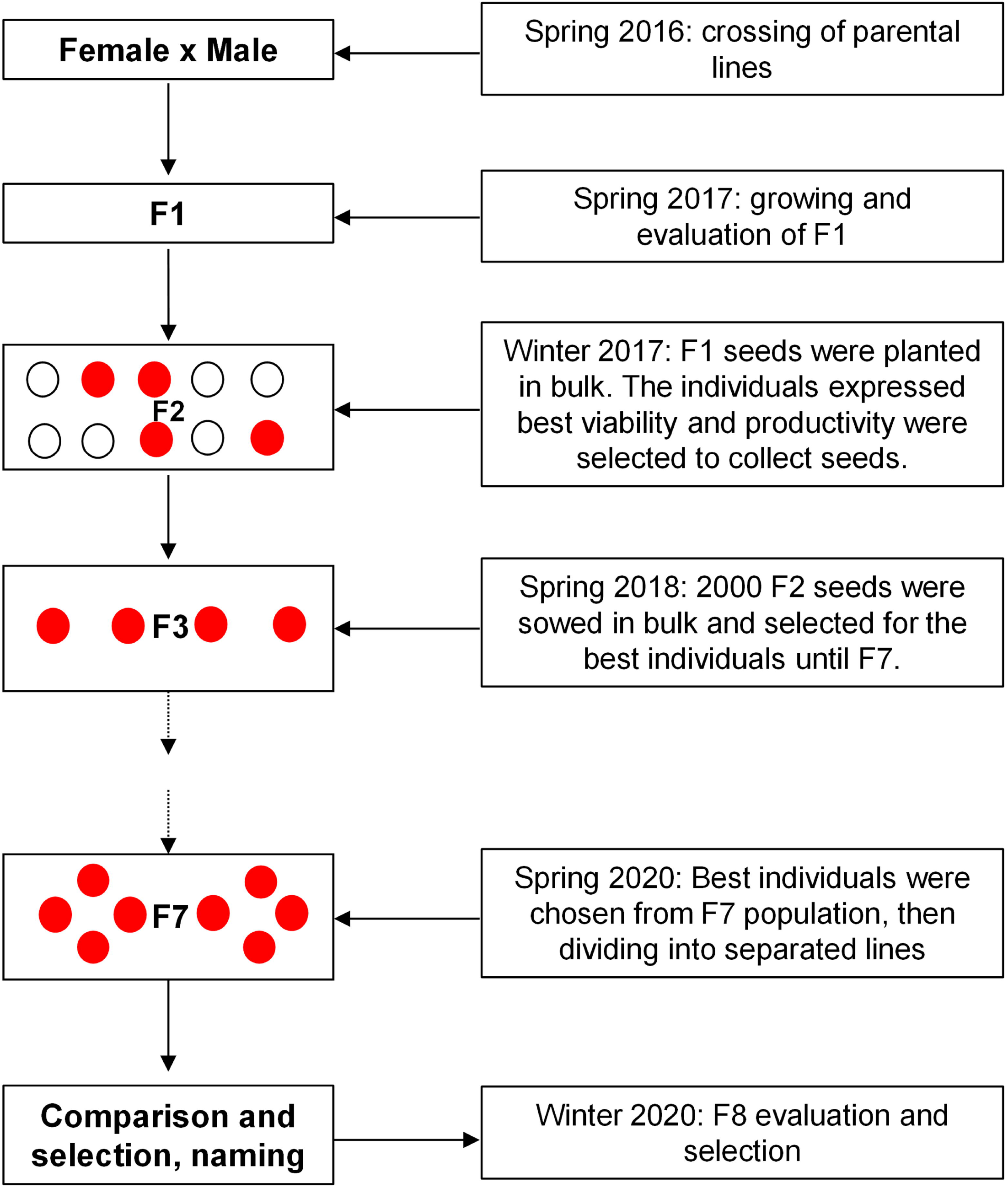
Assessment of Ty QTLs presence in F8 lines. DNA was extracted from the young leaves of at least 4 plants belonging to each line to use as templates for PCR. Plant numbers and line names are shown on top of the panels. Allele name and GAPDH internal control are depicted on the left side of each panel. For assessment of the *Ty-1* alleles, DNA fragments were amplified using CAPS1 forward and reverse primers, then PCR products were digested by *Taq-1* restriction enzyme. Middle lane: *Ty-3* genotype. PCR was conducted using P6-G25 forward and reverse primers. GAPDH (lower lane) was used as the internal control.

Conclusively, the F8 population contains significant good phenotypes which benefit plant health, fruit production and market, such as early harvest (ET4, ET7, and ET10), big fruits (ET1, ET5, ET9), high yield (ET1, ET8, ET10), TYLCV resistance loci (ET3), jointless pedicels (ET2), and partial parthenocarpy (ET9).

## 4 Discussion

Elite tomato lines were developed by crossing between accessions obtained from AVRDC with the Vietnamese domestic cultivars to inherit the advantageous traits while maintaining adaptability to Vietnamese climate conditions. The individual accessions provided by AVRDC possess desired traits, such as high yield and disease resistance. The favorable characteristics of the offspring were re-evaluated in domestic conditions by phenotypic observation or molecular techniques. The selected mother lines were Vietnamese inbred cultivars with high adaptability to climate change, mainly heat resistance. We applied the conventional breeding techniques to develop elite lines with improved yield and different sets of adventitious characteristics that could positively contribute to the gene pool of Vietnamese tomatoes. Since tomato is a self-pollinated species, new cultivars can be derived from a single plant after crossing or by a mixture of offspring (F1). Elite tomato lines were selected from a large pool of progenies based on morphological features, then bulk population breeding was applied to simultaneously maintain the initial variability and eliminate the susceptible individuals (Li and Xu 2022). Therefore, selected ET lines could adapt well to the local conditions.

Plant breeding goals vary depending on the demands of end-users (Acquaah 2016). The farmers would want varieties with high yields, resistance against biotic and abiotic stresses, and early maturing. The processors need to use tomato fruits as raw materials to manufacture industrial products, such as juice and paste, thus, demand deep red color, fewer seeds, high levels of soluble solids, titratable acidity, and consistency (Barrett et al. 2007). On the other hand, consumers favor good taste with increased sweetness, high nutrition, long shelf-life, and beautiful appearance (Acquaah 2016). Several of these crucial characteristics depend on the plant architecture, which includes determinate, semi-determinate and indeterminate growth habits. Determinate varieties stop growing after a certain period, with a flower cluster at the end of each stem. Determinate plants (as observed in the case of ET4, 5, 6, 7, 8, 9, 10) are short and bushy, while semi-determinates, which are similar in shape to determinates, are taller. Indeterminate varieties keep growing for extended periods, and continuously produce new leaves and flowers. Bushy tomato varieties generally produce a higher yield, homogenous fruit set, and have shorter growing time, thus could be harvested earlier, require less labor as it does not require stem pruning, and have better stress resistance. On the other hand, indeterminate lines (such as ET1 and ET3) facilitate higher yield and secure fruit production for long periods, save space, and are suitable for greenhouse conditions, as described in previous studies (McGarry and Ayre 2012; Vicente et al. 2015; Maboko et al. 2017). -Depending on the breeding goals, determinate (ET4, 5, 6, 7, 8, 9, 10), indeterminate (ET1, ET3), or semi-determinate (ET2) lines could be selected as breeding materials in future research. ET3 showed significantly slower fruit ripening and flowering, a typical feature of indeterminate tomato varieties. Slow-ripening and continuous harvest are advantageous to provide fruits for an extended period and off-season. Therefore, it could be adventitious for farmers due to the improved market value. Fruits could be harvested earliest in the determinate lines such as ET7, ET10, and ET4. Early flowering is expected to achieve early production, thus increasing the market price, and reducing production costs. The data in Table 2 proves the correlation between plant growth habits and ripening time.

Productivity is a polygenic trait that directly affects fruit set efficiency and fruit size. In the F8 populations, high-yield lines, such as ET8 and ET10, showed high fruit formation (approximately 120 fruits per plant, **Table 3**). Bigger size fruits of ET1 and ET9 also help to increase productivity, although the difference was not significant in comparison with most of the other lines (**Table 2**). Although all the parental lines of the ET population showed high-yield phenotypes (**Table 1**), not all the progenies could inherit this trait. This observation is consistent with the fact that productivity is governed by multiple genetic factors and that fruit set efficiency seems more important than fruit size in achieving higher productivity.

Tomatoes are produced with the highest yield in Europe, where Belgium was the best producer, with an average of 502.42 tons/ha in 2020 (FAO 2020). However, on average, the European Union only produced 70.91 tons/ha on average, a little lower than Oceania countries which were 73.44 tons/ha. The tomato yield of Asia countries in 2020 was 43.60 tons/ha, which is significantly lower than Europe, Oceania, and America (67.64 tons/ha). This is a big increase compared with 14.19 tons/ha in 1961, however, it still shows a big gap with the developed countries. In Asia, China performed best, producing 58.48 tons/ha. In 2019, Vietnam produced 673,195.5 tons of tomato in a total area of 23,791 ha, giving an average yield of 28.3 tons/ha (Doan et al., 2021). The control C155 variety, which is widely grown in Vietnam in the recent years, produced approximately 34 -43 ton/ha (**Table 3**; Doan et al. 2021; Nguyen et al. 2007). Compared to this production, the ET lines in this study would contribute positively to improving the overall yield of the country, especially ET8 (156.61 tons/ha), ET10 (129.55 tons/ha), and ET1 (114.92 tons/ha).

The TYLCV resistance is important for tomato production worldwide. *Ty-1* and *Ty-3* have been proven to contribute to geminivirus tolerance (Verlaan et al. 2013; Prasad et al. 2020). Plants possessing one or two alleles acquire a significant tolerance to TYLCV infection. These alleles encode for a member of RNA-dependent RNA polymerase and may function by generating siRNAs that support geminivirus DNA silencing through direct methylation. Therefore, ET3 line harboring *Ty-1* and *Ty-3* loci could provide a possible tolerance to TYLCV infection and thus can be an important breeding material for tomato breeders (**Figure 4**).

ET2 showed a jointless phenotype which can confer fruit abscission to protect productivity (**Figure 2**). Abscission benefits plants by removing damaged organs and releasing ripped fruits from stalks, but it also causes the productivity lost when the weather conditions are unfavorable. The jointless trait is desirable in tomato since it helps plants to keep fruits for a longer time and increase the fruit load that each senescent could carry, therefore, highly desired by commercial tomato production.

The parthenocarpy trait of ET9 demonstrated that the mutants toward seed formation could happen naturally, implicating a chance to mimic the alleles by genetic tools. There are parthenocarpy cultivars – which were reported to carry *pat-2* alleles or a novel *pat-k* gene – that have been used as breeding materials in Japan (Gorguet et al. 2008; Takisawa et al. 2017). Parthenocarpy mutant alleles of *SlTPL1* and *IAA9* regulate seed phenotype via controlling the metabolism of phytohormones such as auxin, cytokinins, and abscisic acid (Wang et al. 2009; Mazzucato et al. 2015; He et al. 2021). These alleles were generated by gene modification via RNAi or CRISPR/Cas9-mediated gene editing. The sequencing result of ET9 did not show the mutation in *IAA9* exon 1 (data not shown), suggesting that the parthenocarpy phenotype of this line is due to other factors. Because none of the parental lines has this phenotype, it is implicated that the mutation appeared during the crossing and generating the ET lines.

In conclusion, we successfully generated a population of elite tomato lines which carries a set of superior traits such as high yielding, disease resistance possibility, seedlessness, jointless pedicels, and well-adaptability. These lines positively contributed to the gene pool of Asian tomato and would be used as good materials for future breeding.

## Author contribution statement

C.C.N. and J.-Y.K. conceived the idea and designed the experiments. C.C.N. designed experiments, performed experiments, analyzed data, and wrote the manuscript; C.C.N., T.V.V., H.T.V., N.N.T., X.C.N., V.-A.-K.D., and H.N.T. performed the experiments; R.M.S. contributed to the interpretation and draft of the output; R.M.S., T.V.V., and J.-Y.K. revised the manuscript. All authors read and approved the final manuscript.

## Acknowledgements

This work was supported by the Key Research Funding Program of Vietnam National University of Agriculture (T2018-12-06 TĐ), National Research Foundation of Korea (2019H1D3A1A01102938, 2022R1A2C3010331, 2020R1A6A1A03044344, 2021R1A5A8029490), and the Program for New Plant Breeding Techniques (NBT, PJ01686702), Rural Development Administration (RDA), Korea.

## Data availability

All data generated or analyzed during this study are included in this published article. Additional data will be provided upon request.

## Supplementary data

**Supplementary Table S1**: Material lines obtained from AVRDC.

