## Supplementary Table S1 for "Characterization of yield and fruit quality parameters of Vietnamese elite tomato lines generated through phenotypic selection and conventional breeding methods"

**Supplementary Table S1**: Material lines obtained from AVRDC

| Name | Accession number | Doi | Permanent URL | Accession URL |
|---|---|---|---|---|
| NHG 195 | VI059802 | 10.18730/133BQ$ | https://purl.org/germplasm/id/970a714f-02da-4bbc-ba25-cf74156cdc08 | http://seed.worldveg.org/search/view/passport/VI059802 |
| NHG 191 | VI059798 | 10.18730/133BKY | https://purl.org/germplasm/id/fe2c60f3-e08f-4d7f-a511-01aa1c577364 | http://seed.worldveg.org/search/view/passport/VI059798 |
| NHG 192 | VI059799 | 10.18730/133BMZ | https://purl.org/germplasm/id/2fdb9c1a-335c-493a-8d66-3f6074749a8d | http://seed.worldveg.org/search/view/passport/VI059799 |
| NHG 135 | VI059746 | 10.18730/133ADX | https://purl.org/germplasm/id/abf4c43c-0ba7-4f02-93fa-a42c74c70353 | http://seed.worldveg.org/search/view/passport/VI059746 |
| NHG 188 | VI059796 | 10.18730/133BHW | https://purl.org/germplasm/id/033812d7-f24b-4df7-b0e9-342cc9021bf4 | http://seed.worldveg.org/search/view/passport/VI059796 |
| NHG 176 | VI059786 | 10.18730/1336CB | https://purl.org/germplasm/id/a63e3daf-3eb9-4c85-8182-d892b342b8f8 | http://seed.worldveg.org/search/view/passport/VI059786 |
| NHG 31 | VI059648 | 10.18730/1337D7 | https://purl.org/germplasm/id/e42cfcb6-d638-4e21-ba67-bce00e90aac6 | https://genebank.worldveg.org/#/a/e42cfcb6-d638-4e21-ba67-bce00e90aac6 |
